# Bilateral neuromuscular adaptation to acute unilateral resistance exercise in healthy older adults

**DOI:** 10.1101/2025.02.13.638001

**Authors:** Nishadi N. Gamage, Abdulmajeed Altheyab, Yuxiao Guo, Bethan E. Phillips, George M. Opie, John G. Semmler, Philip Atherton, Mathew Piasecki

**Affiliations:** Centre of Metabolism, Ageing & Physiology, MRC-Versus Arthritis Centre for Musculoskeletal Ageing Research, Nottingham NIHR Biomedical Research Centre, University of Nottingham, United Kingdom; Neurophysiology of Human Movement Laboratory, Discipline of Physiology, School of Biomedicine, University of Adelaide, Australia; Department of Occupational Therapy, College of Applied Medical Sciences, King Saud bin Abdulaziz University for Health Sciences, Saudi Arabia; Institute of Sports Medicine and Health, Chengdu Sport University, Chengdu, China

**Keywords:** Cross-education, motor units, electromyography, older adults, resistance exercise

## Abstract

**Introduction:** Resistance exercise (RE) enhances functionality in older adults and has proven effective as a means of cross-education in scenarios of unilateral disuse. However, the extent to which older adults demonstrate cross-limb transfer at the motor unit (MU) level following a single bout of unilateral RE is unclear.

**Methods:** Thirteen healthy older adults (74.9 ± 4.8 years; 5 females) underwent bilateral neuromuscular assessments pre- and post- a single bout of unilateral RE consisting of 3-4 sets of 8-12 repetitions of leg extension of the dominant (exercise) leg, at 75% of 1 repetition maximum, performed to failure. Maximum voluntary contraction (MVC) and force steadiness (FS) were measured. Central and peripheral features of individual MU were recorded using high-density surface electromyography and intramuscular electromyography (HDs/iEMG), during contractions normalised to 25% MVC.

**Results:** Following unilateral RE, MVC reduced in exercise (-14.8%, *p* < 0.001) and control (-6.9%, *p* = 0.003) legs, with reduced FS performance in the exercise leg compared to the control *(p* = 0.002). MU firing rate increased during contractions normalised to 25% baseline MVC in the exercised leg (*p* < 0.05), with no adaptation in the control leg (*p* > 0.05). All iEMG recorded measures of MU potentials remained unchanged in both legs (all *p* > 0.05).

**Conclusion:** Acute unilateral RE leads to bilateral MVC reduction in older males and females, demonstrating the cross-limb transfer effect. However, adaptation of MU features was only apparent in the exercised limb, and mechanisms underlying the force decline in the non-exercised limb remain uncertain.

## INTRODUCTION

Ageing is associated with a substantial decline in muscle mass, strength, and function leading to detrimental outcomes such as mobility disorders, frailty, increased fall rates, and reduction in general health status (Wilkinson et al., 2018). The age-related decline in muscle function is partly explained by an increase in motor unit (MU) size and reduced MU number (Piasecki et al., 2018), and the consequential reduced control of muscles (Enoka & Duchateau, 2017). Although the age-related loss of motor neurons and muscle fibres are irreplaceable, the structure and function of the musculoskeletal system can be improved through exercise training (McPhee et al., 2016).

Cross-education (CE) describes the widely reported effect of unilateral training conferring benefit to the non-exercised contralateral limb (Altheyab et al., 2024). This is effective in both the upper and lower limbs (Gabriel et al., 2006), following stroke (Dehno et al., 2021, Dragert and Zehr, 2013) and in orthopaedic conditions (Manca et al., 2021), making it particularly relevant for older adults. Although the exact mechanism of cross-limb transfer remains uncertain, it likely has cortical origins (Calvert and Carson, 2022, Ruddy and Carson, 2013).

Exercise-induced decline in force and/or power generation due to impaired muscle activation and/or contractile function is termed performance fatigue and originates from central and peripheral sources (Enoka and Duchateau, 2016). Similar to CE, cross-limb transfer of muscle fatigue is defined as a temporary deficit in performance of the non-exercised contralateral limb following a unilateral fatiguing protocol (Doix et al., 2013, Martin and Rattey, 2007). In young adults, higher-intensity fatiguing contractions are reported to generate greater non-local muscle fatigue effects compared to lower-intensity contractions, particularly in lower limbs (Kawamoto et al., 2014).

Ageing is not a limiting factor in some aspects of cross-limb transfer (Barss et al., 2016). However, there are several knowledge gaps regarding cross-limb transfer, including a lack of data characterising individual MU adaptations in the knee extensors (KE). Additionally, research on this phenomenon in older populations—where cross-limb transfer could be particularly valuable as an intervention—is limited, and to our knowledge, there is no available data following a single bout of unilateral KE fatiguing exercise in older individuals. Therefore, the current study aimed to evaluate the cross-limb transfer effects of acute resistance exercise (RE) on neuromuscular function, and central and peripheral features of MU adaptation in older adults. We hypothesized that both the exercised and control limbs would display reduced function, with MU firing properties altered in both, and peripheral MU features altered in the exercised limb only.

## METHODS

### Ethical approval

This study was approved by the University of Nottingham Faculty of Medicine and Health Sciences Research Ethics Committee (*FHMS - 390-1121*) and was conducted in accordance with the *Declaration of Helsinki*, except for registration in a database. All participants provided written informed consent.

Seventeen healthy older adults (9 females) with a mean age (± SD) of 73.8 (± 4.9) years were enrolled in this study. Participants were excluded if they had a BMI < 18 or > 35 kg/m^2^, any diagnosed neurological, musculoskeletal, cardiovascular (such as uncontrolled hypertension, recent cardiac event), metabolic disorders (such as diabetes), chronic kidney disease, malignancy (in previous 6 months), recent steroid treatment (within 6 months), respiratory diseases or if they engaged in any structured exercise training within the past 6 months. Participants were recreationally active but not involved in standard exercise programs. All participants were advised to refrain from strenuous exercises at least 48 hours before the laboratory visits. Of the 17 participants who completed the study, 13 participants (74.9 ± 4.8 years; 5 females) had sufficient data quality for HDsEMG analyses, therefore data are presented for these only. Of these 13, 11 participants (4 females) were included in iEMG analyses due to the low yield of MUs at one or more time points.

### Screening visit

Prior to study enrolment, participants completed a comprehensive medical screening visit that included blood tests, electrocardiogram, blood pressure assessment, and a health questionnaire which allowed the exclusion of participants against pre-determined criteria as listed above. Additionally, VL muscle ultrasound and 1 repetition maximum (1 RM) assessments were also performed during this visit.

### 1 repetition maximum assessment

The 1 RM is defined as the maximum weight that could be lifted through the prescribed range once only (McNair et al., 2011). A seated leg extension machine was used to measure 1 RM with an initial warm-up of 5-10 low-load repetitions performed on the dominant (right in all) limb. After one minute of rest, participants were asked to perform full knee extension movement with the load at ∼80% of estimated 1 RM. After each successful attempt, the weight was increased progressively until a failed attempt occurred (Seo et al., 2012). Each attempt was separated by one minute rest and 1 RM was obtained within five attempts. The 1 RM was used to determine the load (75% of 1 RM) for the acute RE protocol.

### Muscle ultrasound recording

The cross-sectional area (CSA) of VL muscle of the exercise limb was measured at baseline using an ultrasound probe (LA523 probe, B- mode, frequency range 26- 32 Hz and MyLabTM50 scanner, Esaote, Genoa, Italy) positioned at the anatomical mid-point of VL, which was measured and identified between the greater trochanter and the midline of the patella. A conductive gel was applied to the surface of the probe to improve the fidelity of images. The ultrasound probe was positioned at the VL midpoint and moved in the medial-to-lateral direction to locate the medial and proximal borders of VL where the aponeurosis of VL intersected with the Vastus Intermedius muscle (Guo et al., 2024). Three axial plane images were obtained and the mean area of three images was considered as CSA.

### Study day assessment

Participants arrived at the laboratory at ∼09.00 h (± 1 h) after an overnight fast. The study day assessment included tasks before and immediately after the unilateral RE. All described measures were individually performed on each leg.

### Resistance exercise (RE)

The acute unilateral RE included 3-4 sets of 8-12 repetitions of unilateral leg extension at 75% of their 1 RM, performed to failure. The RE was performed by the dominant limb on a standard leg extension machine with the opposing limb serving as a control. The Borg RPE (rating of perceived exertion) scale was used to determine the participant’s perceived fatigue level (Borg and Kaijser, 2006) and all participants reported a Borg scale of 19-20 (extremely hard - maximal exertion) following the final set.

### Functional properties

#### Muscle strength

A purpose-built isometric dynamometer (Load cell amplifier; LCA1,12V1A medical PSU, GDM25B12-P1J, Sunpower Electronics, Reading, United Kingdom) was used to measure MVC for knee extension. Participants were seated with their hip and knee flexed at approximately 90° and a seat belt was used to avoid hip or trunk movement during the leg extension. The leg to be tested was securely strapped to a plate connected to a force transducer at the ankle. After three moderate-intensity warm-ups, participants were instructed to perform three maximal isometric knee extensions with a 60-second rest between trials. Real-time visual feedback (Spike software v9.06; Cambridge Electronic Design, Cambridge, United Kingdom) and verbal encouragement were provided throughout all attempts.

#### High-density surface electromyography (HDsEMG)

A semi-disposable high-density surface electromyography (HDsEMG) array (64 electrodes, 13 × 5, 8 mm, I.E.D., GR08MM1305, OT Bioelettronica, Inc., Turin, Italy) was positioned over the VL muscle belly with approximate orientation of the muscle fascicles (proximal to distal). Prior to this, the skin was prepared by shaving, applying light abrasion, and cleansing with 70% ethanol. These HDsEMG electrodes were secured using flexible tape and remained in place during the intervention and post-RE assessments. A ground electrode (WS2, OTBioelettronica, Turin, Italy) was positioned around the ankle of the tested leg. HDsEMG signals were acquired in a monopolar configuration, amplified (× 256) with filtering set at 10–500 Hz and digitally converted at 2000 Hz by a 16-bit wireless amplifier (Sessantaquattro, OTBioelettronica, Turin, Italy) and transferred to a PC for offline analysis.

### Intramuscular EMG (iEMG)

#### Motor point identification

The motor point of VL was identified using low-intensity percutaneous electrical stimulations (400V, pulse width 50 μS, current ∼10 mA; delivered via a Digitimer DS7A, Welwyn Garden City, United Kingdom). The site of VL which produced the largest visible twitch from the smallest electrical current was considered the motor point and its proper localization is considered critical for electrode positioning (Botter et al., 2011) .

#### Force and electromyography recording

Intramuscular electromyography (iEMG) recordings were obtained using a disposable concentric needle (model N53153, Teca, Hawthorne, New York, United States) with a recording area of 0.07 mm^2^. The grounding electrode was placed over the patella of the tested leg. Subsequently, the surrounding skin at the motor point was prepared by shaving and cleansing using an alcohol adhesive wipe. Participants were instructed to relax their tested leg to enable insertion of the needle into the muscle belly of VL until the appropriate depth of 1.0- 2.5 cm was reached. Next, participants were instructed to perform isometric leg extension, and the signal quality was inspected, and the needle was adjusted where necessary. Subsequently, the needle was repositioned by rotating the bevel 180° and slightly retracted by ∼5 mm to ensure that different depths of the muscle were captured (Jones et al., 2021). Once the appropriate contraction sequence (sets of 25% of MVC) was completed (minimum of four recordings from distinct perspectives/regions), the needle was withdrawn and disposed of. The EMG signals were recorded at 50 kHz and bandpass filtered at 10 Hz to 10 kHz (CED Micro 1401; Cambridge Electronic Design, Cambridge, United Kingdom) and visualised using Spike2 (version 9, CED) software to provide real-time visual feedback on a screen positioned in front of the participants.

### Data analysis

#### Neuromuscular function

The highest isometric knee extension of the three recorded values was considered the MVC. To quantify the force steadiness (FS), force signals during the plateau phase of a sustained voluntary contraction performed at 25% MVC were low-pass filtered at 20 Hz (fourth-order Butterworth digital filter), and the coefficient of variation (CoV) of force was quantified. Ultrasound images were analysed using ImageJ software (National Institutes of Health, United States) and the average of three ultrasound images was considered as VL CSA of the exercise limb.

#### HDsEMG analysis

The HDsEMG signals recorded during contractions held at 25% MVC were analysed offline using the convolution kernel compensation method (Holobar and Zazula, 2007). MU filters and corresponding spike trains estimated with this process (Francic and Holobar, 2021) were visually inspected and optimised using custom software (DEMUSE, MATLAB) and standardised procedures (Del Vecchio et al., 2020). Only MUs with a pulse-to-noise ratio (PNR) ≥ 30 dB were included for further analyses. After editing each MU spike train, MUs were tracked from pre- to post-intervention via concatenation of signals and application of MU filters from one contraction (within each leg) to the other.

MU firing rate (MUFR) at the recruitment and derecruitment phases (Figure 1D) of isometric knee extension was calculated as the mean instantaneous firing rate (FR) of the first 5 observations of each MU. MUFR during the sustained phase was the mean instantaneous FR throughout this segment. FR variability was considered as the CoV of the inter-spike interval (ISI) during the plateau phase. MU recruitment and derecruitment threshold was defined as the force level corresponding to the first and last observed firing of each MU, respectively. MUs that were tracked from pre- to post-intervention were used to compute a cumulative spike train (CST), which represents the neural drive to the muscle (Martinez-Valdes et al., 2022).

**Figure 1.**
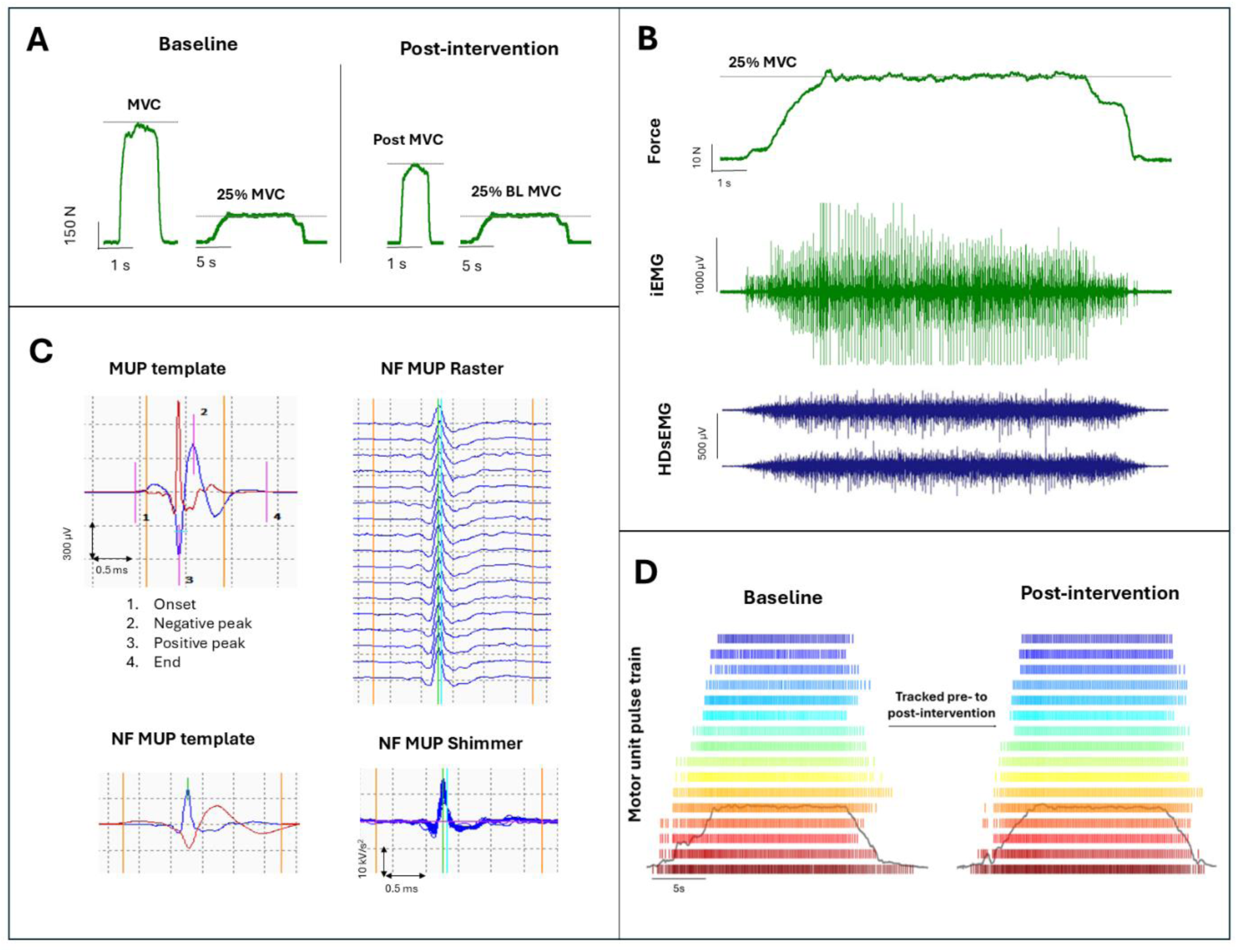
Example knee extensor force traces of an older male participant showing MVC and 25% MVC of the exercise leg obtained at baseline and post-RE intervention (A). Example force trace (upper) with corresponding intramuscular electromyography (iEMG) trace (middle) and a subset of 2 HDsEMG channels (lower) obtained during a sustained isometric knee extension at 25% of MVC (B). Example iEMG recorded MUP and corresponding near fibre (NF) MUP sampled with iEMG, including consecutive NFMUPs shown as raster and shimmer plots (C). Example HDsEMG recorded MU pulse trains (firing times) of MUs tracked from baseline to post-intervention. Grey line indicates force trace with plateau held at 25% MVC (D). *Abbreviations*: *MVC, maximal voluntary contraction; BL, baseline; MUP, motor unit potential; NF, near fibre*.

#### iEMG analysis

Decomposition and iEMG analysis were performed using Decomposition-based quantitative electromyography (DQEMG) software. EMG signal decomposition involves the process of breaking down a complex EMG signal into its motor unit potential trains (MUPTs) (Parsaei et al., 2009) which includes preprocessing of signal, signal segmentation and MUP detection, clustering the identified MUPs, and supervised classification of detected MUPs (Stashuk, 2001). Visual inspection of individual MUP templates was performed to ensure cursors were positioned accurately at the onset, endpoint, positive peak, and negative peak of the waveforms (Figure 1C). Where required, cursors were adjusted and repositioned during visual inspection. MUPTs were excluded if they contained MUPs from multiple MUs or fewer than 40 MUPs (Inns et al., 2022).

MUP area, measured in µVms, was defined as the integral of the absolute value of the MUP values between the onset and end markers, multiplied by the sampling time interval (Piasecki et al., 2021). MUP complexity was assessed using the number of turns in the MUP template and turns were defined as the number of significant changes in direction within the MUP duration (height > 20 µv) (Piasecki et al., 2021). The MUP negative peak ratio was calculated based on the absolute value of the ‘rise’ of the MUP template negative peak (within the 500 μs interval before the negative peak), divided by the ‘fall’ of the MUP template negative peak (within the 500 μs interval after the negative peak) (Jones et al., 2023). Individual MUP templates were used to calculate a near-fibre MUP (NFM) (Piasecki et al., 2021). Individual NFMs were visually inspected using DQEMG software, and any that were contaminated from other NFMs were excluded from the analysis. NFM parameters were used to estimate NF jiggle as an estimate of neuromuscular junction transmission instability (Piasecki et al., 2021).

### Statistical analysis

All statistical analysis was conducted using RStudio (Version 4.4.0). To assess differences in MVC, FS, HDsEMG, and iEMG MU variables, separate linear mixed-effects models were constructed with fixed effects of Time and Leg, and their interaction (Leg × Time). The model included random intercepts for each Subject, with Leg nested within Subject [e.g. (1 | Subject/Leg)] to account for repeated measures. For tracked MU models, MUs were also nested within Subject. All linear-mixed effects models were generated using *lmer* R package (Bates et al., 2015). Where significant main effects were detected, pairwise *post-hoc* tests of the estimated marginal means (EMMs) were performed (Lenth and Lenth, 2018). Delta CST values were compared across limbs with a repeated measures t-test. Pearson correlation test was used to assess the strength, direction, and significance of the relationship between delta MVC and delta FR. The tabulated results are displayed as EMMs, 95% confidence intervals (CI), and *p* values. Statistical significance was accepted at *p* < 0.05.

## RESULTS

### Participant characteristics

Participant characteristics are shown in Table 1. All participants completed the experimental session without any adverse effects.

**Table 1.** Descriptive characteristics of participants *(n = 13; 5 female)*

|  |  |
| --- | --- |
| Age (years) | 74.9 (4.8) |
| Height (cm) | 167.6 (6.8) |
| Weight (kg) | 73.5 (11.7) |
| BMI (kg/m<sup>2</sup>) | 26.1 (3.0) |
| VL CSA (cm<sup>2</sup>) | 17.1 (4.3) |

Data shown as mean (standard deviation). VL CSA obtained from the exercise (dominant) leg only at baseline. *Abbreviations: BMI, body mass index; VL CSA, vastus lateralis cross-sectional area.*

### Functional properties

All model outputs including estimated marginal means (EMM) for functional properties are shown in Table 2. Knee extensor MVC at each timepoint (pre and post) for the exercise (right) and control (left) legs for 13 participants are shown in Figure 2A. There was a significant leg x time interaction on knee extensor MVC (*p* = 0.013), and a main effect of time (*p* = 0.003), with no main effect of leg (*p* = 0.388). EMMs show a mean decrease in MVC of 14.8% (*p* < 0.001) in the exercise leg, and of 6.9% in the control leg (*p* = 0.003).

**Figure 2.**
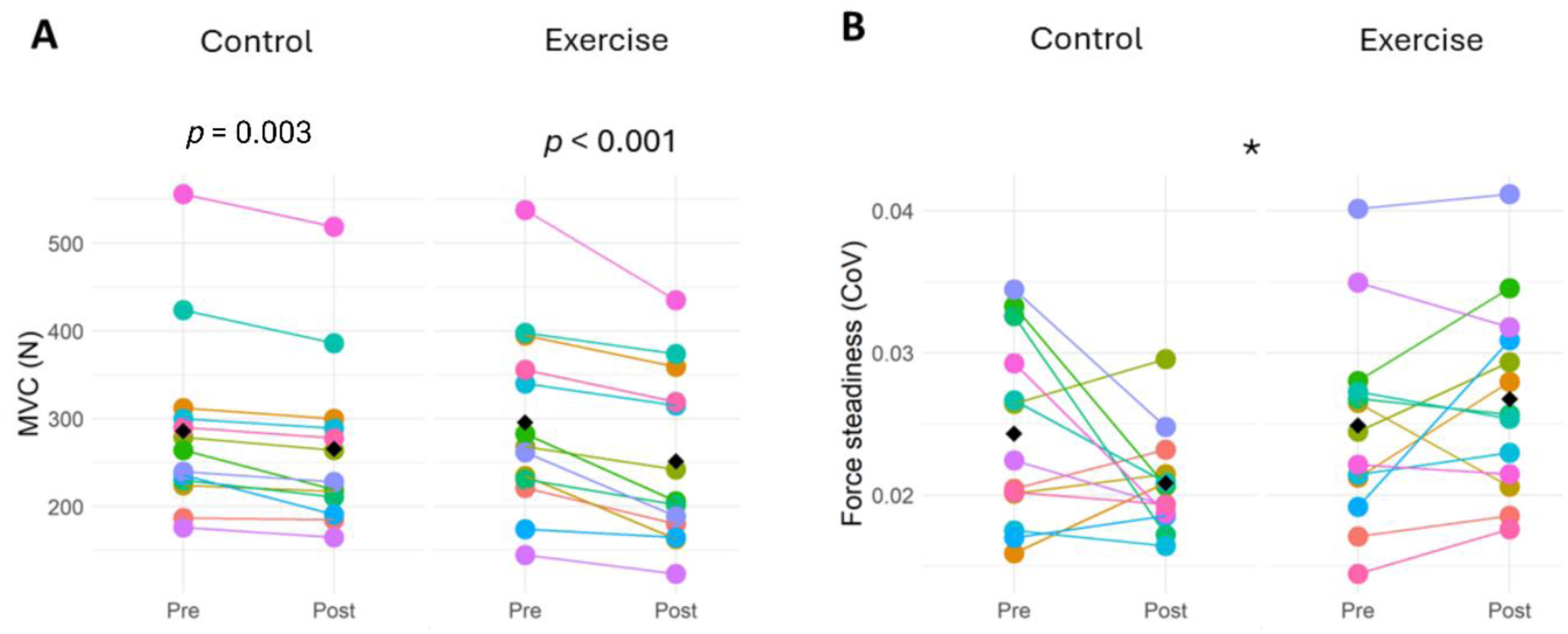
Effect of acute unilateral RE on knee extensor MVC (A) and force steadiness (B) in control (left) and exercise (right) legs. Data points are colour-coded for individual participants with the mean of means indicated by black diamonds. *Significant difference between the exercise and control legs at post-intervention. *Abbreviations*: *MVC, maximal voluntary contraction; CoV, Coefficient of variation of force at 25% of MVC*.

**Table 2.** Summary of linear regression analyses for functional properties and HDsEMG non-tracked MUs.

|  | Pre-intervention |  | Post-intervention |  | Leg | Time | Leg x Time |
| --- | --- | --- | --- | --- | --- | --- | --- |
|  | Control | Exercise | Control | Exercise |  |  |  |
| ***Functional properties*** |  |  |  |  |  |  |  |
| MVC (N) | 286<br>(226 – 346) | 296<br>(235 – 356) | 266<br>(205 – 326) | 252<br>(191 – 312) | 0.388 | 0.003* | 0.013* |
| Force steadiness (CoV) | 0.024<br>(0.020 – 0.028) | 0.025<br>(0.021 – 0.029) | 0.021<br>(0.018 – 0.024) | 0.027<br>(0.024 – 0.030) | 0.742 | 0.062 | 0.032* |
| ***HDsEMG non-tracked MU*** |  |  |  |  |  |  |  |
| No of MUs | 177 | 178 | 133 | 128 |  |  |  |
| FR recruitment (pps) | 6.21<br>(5.76 – 6.66) | 5.89<br>(5.44 – 6.34) | 6.01<br>(5.55 – 6.48) | 6.69<br>(6.23 – 7.16) | 0.001* | 0.182 | 0.001* |
| FR plateau (pps) | 7.43<br>(6.75 – 8.11) | 7.32<br>(6.65 – 8.00) | 7.26<br>(6.57 – 7.95) | 8.66<br>(7.97 – 9.35) | 0.001* | 0.280 | 0.001* |
| FR derecruitment (pps) | 5.04<br>(4.62 – 5.46) | 5.09<br>(4.67 – 5.51) | 5.12<br>(4.69 – 5.55) | 6.07<br>(5.64 – 6.50) | 0.001* | 0.467 | 0.001* |
| FR variability (%) | 0.141<br>(0.127 – 0.155) | 0.124<br>(0.110 – 0.138) | 0.137<br>(0.121 – 0.153) | 0.112<br>(0.096 – 0.128) | 0.607 | 0.619 | 0.476 |
| Recruitment threshold (N) | 33.3<br>(24.8 – 41.9) | 33.6<br>(25.1 – 42.2) | 35.6<br>(26.9 – 44.3) | 31.8<br>(23.1 – 40.5) | 0.331 | 0.277 | 0.176 |
| Derecruitment threshold (N) | 30.9<br>(22.8 – 39.0) | 34.5<br>(26.4 – 42.6) | 31.1<br>(22.9 – 39.3) | 35.3<br>(27.1 – 43.5) | 0.441 | 0.906 | 0.810 |

Data presented as estimated marginal means (lower 95% CI, upper 95% CI). Final three columns show *p* values. Abbreviations: *MVC*, maximal voluntary contraction; *CoV*, Coefficient of variation of force at 25% of MVC; *FR*, firing rate. \*Statistically significant ( $p < 0.05$ ).

For FS, there was a significant leg x time interaction (*p* = 0.032), but no effect of leg (*p* = 0.742) or time (*p* = 0.062). *Post-hoc* analysis indicated that FS CoV was comparable between the legs at baseline (*p* = 0.743, however, following the intervention, FS CoV was higher in the exercise leg compared to the control leg *(p* = 0.002) (Figure 2B).

### Central motor unit characteristics

From HDsEMG, a total of 616 MUs from 13 participants were sampled in both limbs at both timepoints. Individual MU counts and all model outputs are shown in Table 2.

MU firing rate (MUFR) at recruitment showed a significant leg x time interaction (*p* = 0.001), a significant effect of leg (*p* = 0.001), but no effect of time (*p* = 0.182). *Post-hoc* analysis showed that, at baseline, MUFR at recruitment was greater in the control leg compared to the exercise leg (*p* = 0.02), however post-intervention, it was greater in the exercise leg (*p* < 0.001). Moreover, MUFR at recruitment did not differ in the control leg from pre- to post-intervention (*p* = 0.182), but for the exercise leg, MUFR at recruitment was significantly greater at post compared to baseline (*p* < 0.001) (Table 2, Figure 3A).

**Figure 3.**
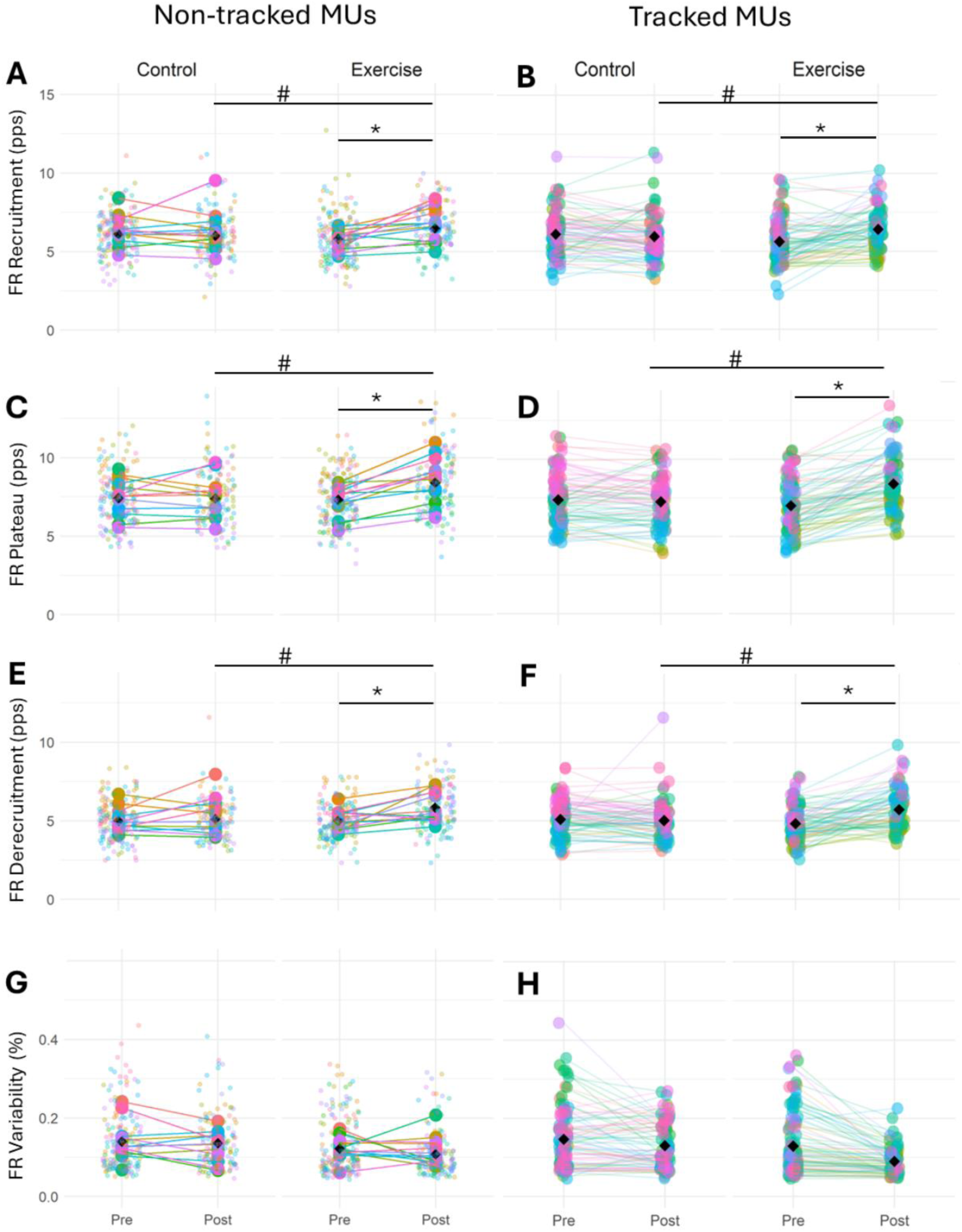
Motor unit firing rate (MUFR) at 25% MVC for non-tracked and tracked motor units in the exercise and control legs before and after the unilateral RE intervention. MUFR at the recruitment phase for non-tracked (A) and tracked (B) motor units. MUFR at plateau phase for non-tracked (C) and tracked (D) motor units. MUFR at derecruitment phase for non-tracked (E) and tracked (F) motor units. MUFR variability for non-tracked (G) and tracked (H) motor units during the plateau phase. Data points are colour-coded for individual participants with the mean of means indicated by black diamonds. Statistical analyses are based on multi-level mixed effects linear models. *Significantly different between pre and post timepoints (*p* < 0.001). ^#^ Significant difference between the exercise and control legs at post intervention (*p* < 0.001). *Abbreviations: FR, firing rate; MUs, motor units*.

During the plateau phase, there was a significant leg x time interaction (*p* < 0.001) and a significant effect of leg (*p* < 0.001), but there was no main effect of time (*p* = 0.280). While MUFR at plateau was similar between the legs at baseline (*p* = 0.477), following the intervention, FR at plateau was greater for the exercise leg compared to the control leg (*p* < 0.001). Additionally, while MUFR at plateau did not change in the control leg across timepoints, it was significantly greater following the intervention compared to the baseline for the exercise leg *(p* < 0.001) (Table 2, Figure 3C).

For MUFR at derecruitment, there was a significant leg x time interaction (*p* < 0.001) and a significant effect of leg (*p* < 0.001), but there was no significant effect of time (*p* = 0.467). *Post-hoc* analysis showed that MUFR at derecruitment was comparable between legs at baseline (*p* = 0.622), but following the intervention, MUFR at derecruitment was greater for the exercise leg compared to the control leg (*p* < 0.001). Additionally, although MUFR at derecruitment did not change in the control leg from baseline to post (*p* = 0.466), there was greater MUFR at derecruitment at post compared to baseline for the exercise leg (*p* < 0.001) (Table 2, Figure 3E).

There was no significant leg x time interaction for MUFR variability (*p* = 0.476) and no significant effect of leg (*p* = 0.607), or time (*p* = 0.619) (Table 2, Figure 3G).

For MU recruitment threshold, there was no significant leg x time interaction (*p* = 0.176), or significant effects of leg (*p* = 0.331) or time (*p* = 0.277) (Table 2, Figure 4A). Similarly, for MU derecruitment threshold, there were no significant leg x time interaction (*p* = 0.810), or significant effects of leg (*p* = 0.441), or time (*p* = 0.906) (Table 2, Figure 4C).

**Figure 4.**
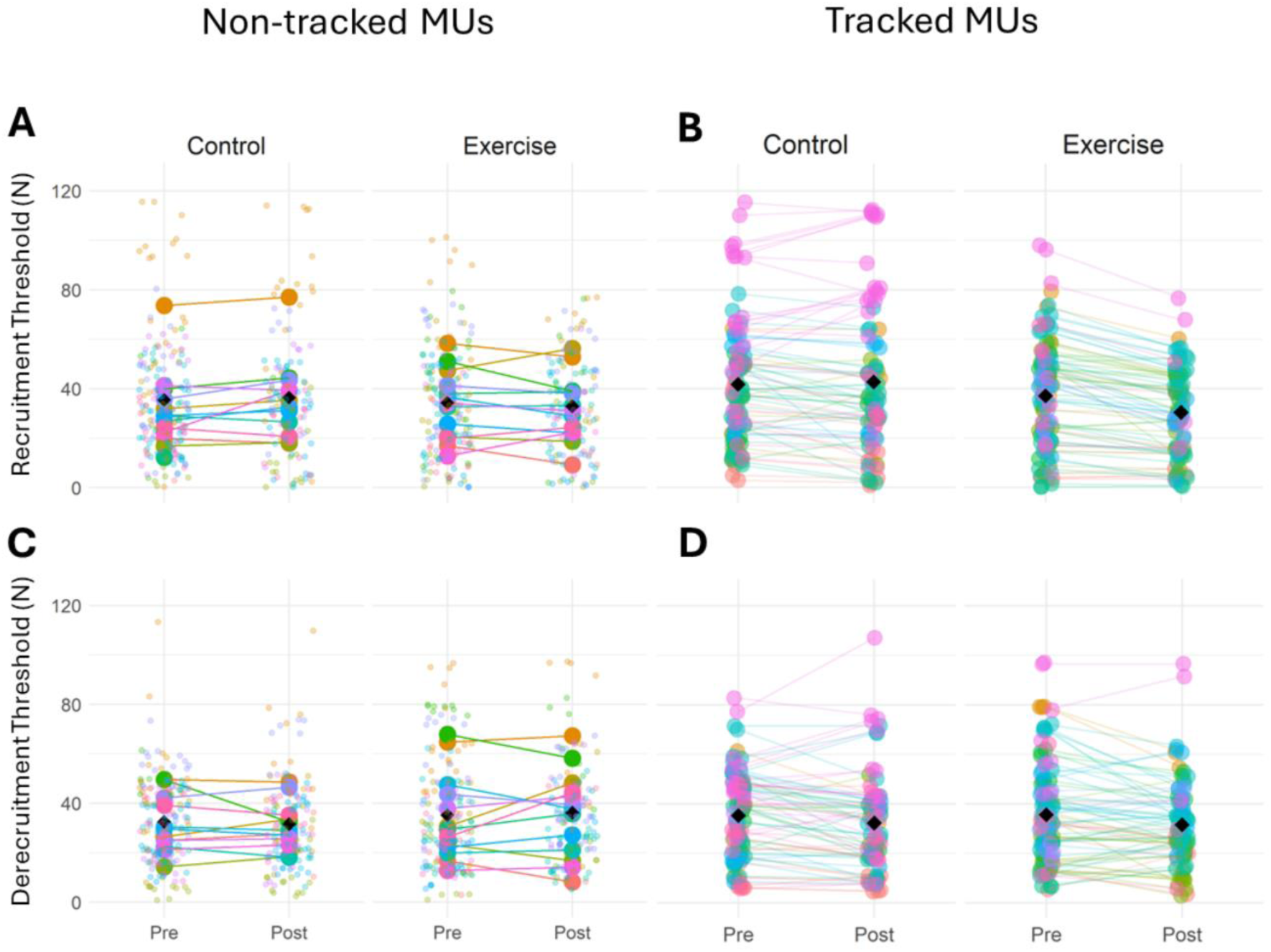
Motor unit recruitment and derecruitment thresholds at 25% MVC for non-tracked and tracked motor units in the exercise and control legs before and after the unilateral RE intervention. MU recruitment threshold for non-tracked (A) and tracked (B) motor units. MU derecruitment threshold for non-tracked (C) and tracked (D) motor units. Data points are colour-coded for individual participants with the mean of means indicated by black diamonds. Statistical analyses are based on multi-level mixed effects linear models.

### Central motor unit characteristics in tracked MUs

A total of 374 MUs from 13 participants were successfully tracked from pre- to post-intervention: 179 in the control leg and 195 in the exercised leg. All model outputs for tracked MUs are shown in Table 3.

**Table 3.** Summary of linear regression analyses for HDsEMG tracked MUs.

|  | Pre-intervention |  | Post-intervention |  | Leg | Time | Leg x Time |
| --- | --- | --- | --- | --- | --- | --- | --- |
|  | Control | Exercise | Control | Exercise |  |  |  |
| FR recruitment (pps) | 6.28<br>(5.66 – 6.90) | 5.82<br>(5.21 – 6.43) | 6.17<br>(5.54 – 6.80) | 6.76<br>(6.13 – 7.39) | 0.008* | 0.545 | 0.001* |
| FR plateau (pps) | 7.33<br>(6.57 – 8.08) | 7.12<br>(6.37 – 7.86) | 7.16<br>(6.40 – 7.93) | 8.69<br>(7.93 – 9.46) | 0.257 | 0.408 | 0.001* |
| FR derecruitment (pps) | 5.14<br>(4.65 – 5.63) | 5.00<br>(4.52 – 5.48) | 5.06<br>(4.56 – 5.56) | 6.05<br>(5.55 – 6.55) | 0.303 | 0.576 | 0.001* |
| FR Variability (%) | 0.148<br>(0.130 – 0.168) | 0.133<br>(0.115 – 0.152) | 0.134<br>(0.114 – 0.155) | 0.097<br>(0.077 – 0.117) | 0.095 | 0.152 | 0.110 |
| Recruitment threshold (N) | 39.5<br>(28.9 – 50.2) | 38.5<br>(28.0 – 49.0) | 38.8<br>(28.1 – 49.6) | 33.3<br>(22.6 – 44.1) | 0.688 | 0.800 | 0.222 |
| Derecruitment threshold (N) | 34.7<br>(26.5 – 43.0) | 37.3<br>(29.1 – 45.5) | 31.4<br>(23.0 – 39.9) | 34.7<br>(26.3 – 43.1) | 0.223 | 0.142 | 0.839 |

Data presented as estimated marginal means (lower 95% CI, upper 95% CI). Final three columns show $p$ values. *Abbreviations: FR, firing rate.* \*Statistically significant ( $p < 0.05$ ).

For MUFR at recruitment, there was a significant leg x time interaction (*p* < 0.001) and a significant effect of leg (*p* = 0.008), but there was no effect of time (*p* = 0.545). *Post-hoc* analysis indicated that at baseline, MUFR at recruitment was higher in the control leg compared to the exercise leg (*p* = 0.008), whereas post-intervention, MUFR at recruitment was higher in the exercise leg compared to the control leg (*p* = 0.003). Additionally, MUFR at recruitment did not differ in the control leg from pre to post (*p* = 0.545), but for the exercise leg, it was significantly greater at post compared to the baseline (*p* < 0.001) (Table 3, Figure 3B).

There was a significant leg x time interaction for MUFR at the plateau (*p* < 0.001), but there were no effects of leg (*p* = 0.257) or time (*p* = 0.408). *Post-hoc* analysis showed that MUFR at the plateau was comparable between the legs at baseline (*p* = 0.257), but post-intervention, it was greater in the exercise leg compared to the control leg (*p* < 0.001). Moreover, although MUFR at the plateau did not differ in the control leg across timepoints (*p* = 0.408), MUFR at the plateau significantly increased from baseline to post in the exercise leg (*p* < 0.001) (Table 3, Figure 3D).

Considering MUFR at derecruitment, there was a significant leg x time interaction (*p* < 0.001), but there was no difference between the legs (*p* = 0.303) or timepoints (*p* = 0.510). *Post-hoc* analysis indicated that MUFR at derecruitment did not differ between the legs at baseline (*p* = 0.303), but following the intervention, MUFR at derecruitment was significantly greater for the exercise leg compared to the control leg (*p* < 0.001). Additionally, MUFR at derecruitment in the control leg did not change across timepoints (*p* = 0.576), but for the exercise leg, MUFR at derecruitment was significantly greater at post compared to the baseline (*p* < 0.001) (Table 3, Figure 3F).

Considering FR variability, there was no significant interaction between leg x time (*p* = 0.110) and no significant differences between legs (*p* = 0.095) nor timepoints (*p* = 0.152) (Table 3, Figure 3H).

For the recruitment threshold, there was no leg x time interaction (*p* = 0.222), or significant effects of leg (*p* = 0.688), or time (*p* = 0.800) (Table 3, Figure 4B). Similarly, for the derecruitment threshold, there was no leg x time interaction (*p* = 0.839), or significant effects of leg (*p* = 0.223), or time (*p* = 0.839) (Table 3, Figure 4D).

### Cumulative spike train

MUs that were tracked from pre- to post-intervention were used to compute the cumulative spike train (CST) for each leg at each timepoint. The mean pre to post difference in the CST was greater in the exercised than the control leg (*p* = 0.043), and the same was true of the percentage difference (*p* = 0.022) (Figure 5).

**Figure 5.**
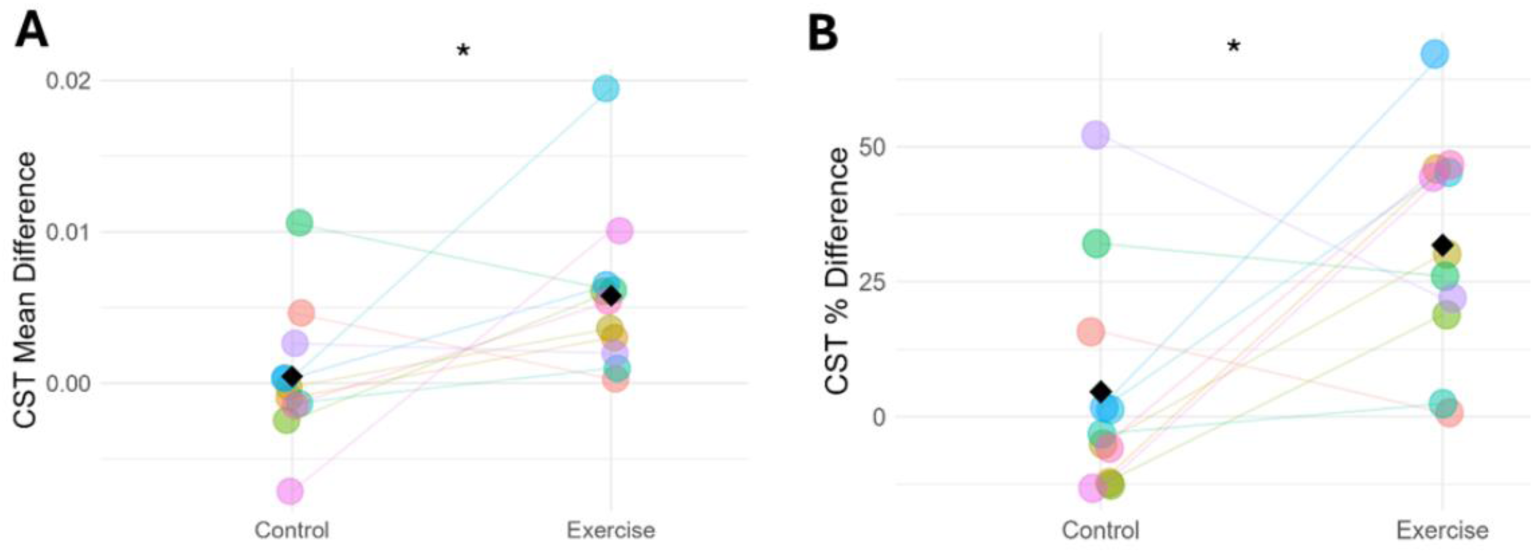
The difference from pre- to post-intervention of the cumulative spike train shown as an absolute value (A) and as a percentage (B). Data points are colour-coded for individual participants with means indicated by black diamonds. *Significantly different between the exercise and control legs (*p* < 0.05). *Abbreviations: CST, cumulative spike train*.

### Correlation between changes in MUFR and MVC

There was a weak yet statistically significant correlation between delta FR and delta MVC in the exercise leg (*r* = -0.566, *p* = 0.043), which was not observed in the control leg (*r* = -0.330, *p* = 0.271) (Figure 6).

**Figure 6.**
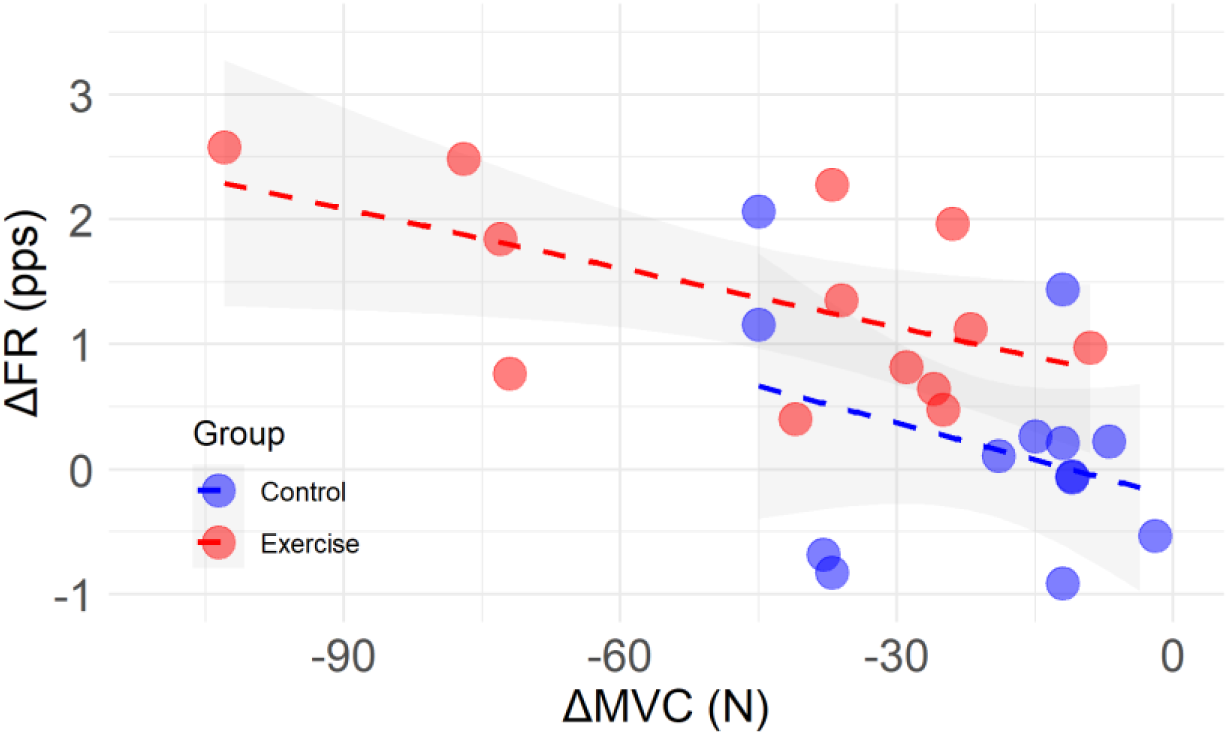
Correlation of change in firing rate as a function of change in MVC, from pre- to post-intervention in the control (blue) and exercise (red) legs. *Abbreviations: FR, firing rate; pps, pulses per second; MVC, maximal voluntary contraction*.

### Motor unit characteristics using iEMG

A total of 841 MUPs were recorded from both legs at both time points for 11 participants (4 females). All Individual MU counts and model outputs for iEMG data are shown in Table 4.

**Table 4.** Summary of linear regression analyses for iEMG motor unit properties.

|  | Pre-intervention |  | Post-intervention |  | Leg | Time | Leg x Time |
| --- | --- | --- | --- | --- | --- | --- | --- |
|  | Control | Exercise | Control | Exercise |  |  |  |
| No of MUPs | 306 | 238 | 142 | 155 |  |  |  |
| Area (μV.ms) | 828<br>(744 - 913) | 901<br>(812 - 991) | 781<br>(675 - 886) | 796<br>(694 - 899) | 0.119 | 0.392 | 0.461 |
| MUP Turns | 4.42<br>(4.03 - 4.82) | 4.53<br>(4.12 - 4.93) | 4.60<br>(4.16 - 5.04) | 4.61<br>(4.17 - 5.05) | 0.503 | 0.332 | 0.714 |
| NF Jiggle (%) | 17.4<br>(14.8 -<br>19.9) | 18.0<br>(15.4 -<br>20.6) | 17.8<br>(15.0 -<br>20.6) | 18.0<br>(15.2 -<br>20.7) | 0.472 | 0.673 | 0.738 |
| Neg Pk ratio | 2.01<br>(1.72 -<br>2.29) | 1.99<br>(1.69 -<br>2.29) | 2.24<br>(1.90 -<br>2.58) | 1.89<br>(1.55 -<br>2.22) | 0.896 | 0.168 | 0.164 |

Data presented as estimated marginal means [lower 95% CI, upper 95% CI]. Final three columns show $p$ values. Abbreviations: MUP, motor unit potential; NF, near fibre.

For MUP area, there were no leg x time interaction (*p* = 0.461), or significant effects of leg (*p* = 0.119), or time (*p* = 0.392). Similarly, for MUP turns there were no significant leg x time interaction (*p* = 0.714) or effects of leg (*p* = 0.503), or time (*p* = 0.332). For NF jiggle, there was no leg x time interaction (*p* = 0.738), or significant effects of leg (*p* = 0.472), or time (*p* = 0.673). Additionally, for the negative peak ratio, there were no significant leg x time interaction (*p* = 0.164), or effects of leg (*p* = 0.896), or time (*p* = 0.168) (Table 4, Figure 7).

**Figure 7.**
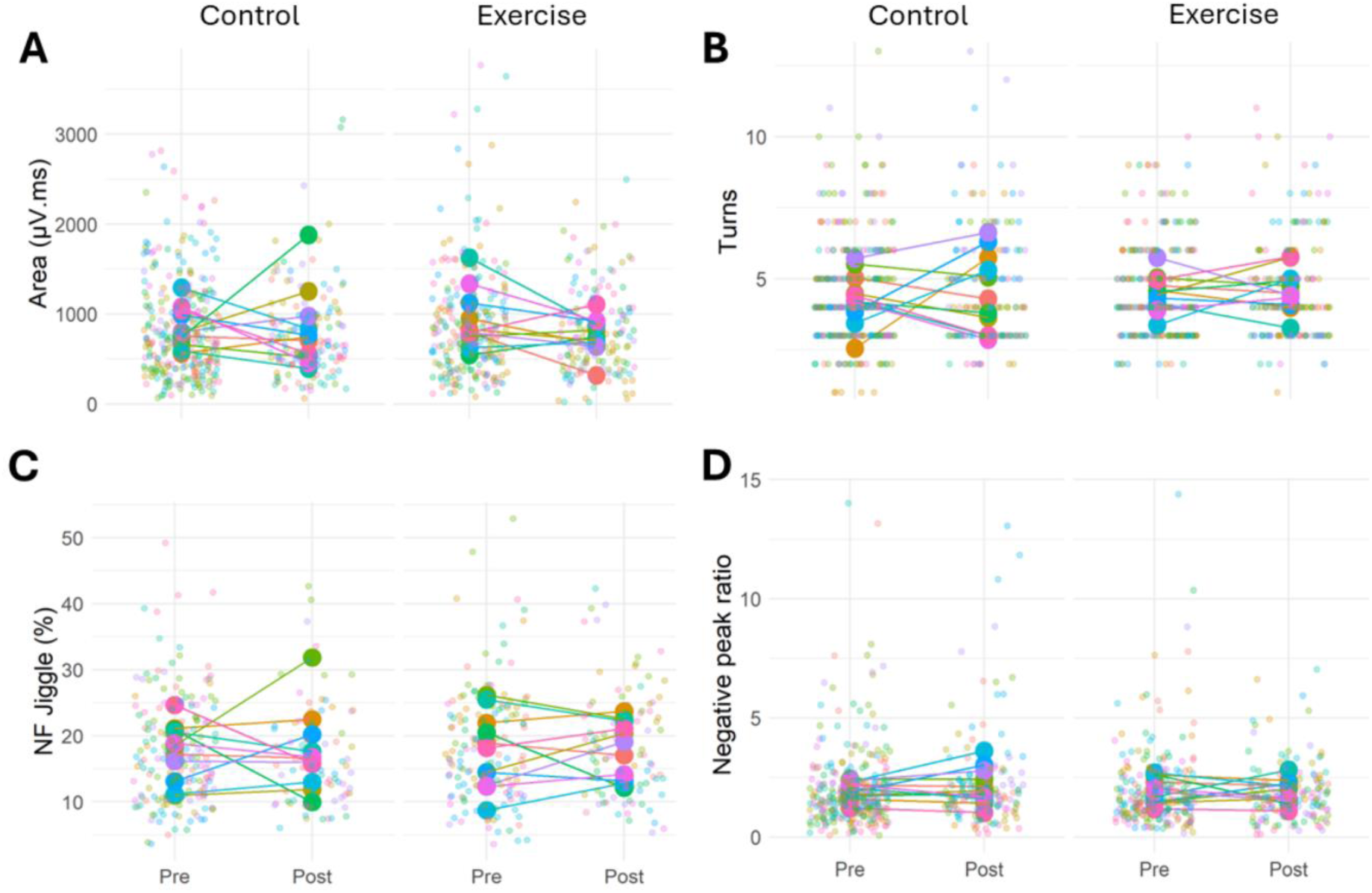
Motor unit characteristics assessed using iEMG. Motor unit Area (A), number of turns (B), neuromuscular junction transmission instability (Near fibre jiggle) (C), and negative peak ratio (D) in the exercise and control legs before and after the unilateral RE intervention. Data points are colour-coded for individual participants. Statistical analyses are based on multi- level mixed effects linear models.

## DISCUSSION

The present study evaluated the effect of acute unilateral knee extensor RE on bilateral neuromuscular function and VL MU features in healthy older adults. The main findings indicate that a single bout of unilateral resistance exercise is sufficient to elicit reductions in maximal knee extensor voluntary force in the exercised and to a lesser extent, in the contralateral non-exercised limb, demonstrating a cross-limb transfer of muscle performance fatigue. However, the majority of markers of individual MU function were altered only in the exercise leg, with MUFR increasing during all phases of a contraction normalised to 25% of baseline MVC, with no notable adaptation in the non-exercise limb.

The resistance-based exercise used in this study was deliberately selected to induce performance fatigue in VL which plays a crucial role in daily activities. Resistance exercise-induced reduction in force of ∼15% in the exercise limb and ∼7% in the non-exercise limb in the current study supports previous findings in young adults (Doix et al., 2013, Halperin et al., 2014, Kawamoto et al., 2014, Martin and Rattey, 2007). Similarly, impaired force accuracy was only apparent in the exercised limb, supportive of previous fatigue-related findings in young adults (Singh et al., 2010).

The decline in contractile function leading to reduced MVC in the exercise leg is undoubtedly partly caused by molecular mechanisms (e.g. reduced sarcoplasmic reticulum Ca^2+^ release, impaired myofibrillar force production, and reduced myofibrillar Ca^2+^ sensitivity (Allen et al., 2008, Debold and Westerblad, 2024)), and the increase in MUFR reported here, which correlated with declines in MVC, likely occurred to compensate for this. Indeed, the loss of force in the exercised leg was 2-fold greater than the control. However, mechanisms explaining the loss in the control leg are less clear.

Earlier studies of cross-limb transfer showed voluntary activation of plantar flexors was reduced following eccentric contractions, in the exercised and contralateral non-exercised limb (Marathamuthu et al., 2022), and separately, sustained fatigue protocols impaired voluntary activation bilaterally following unilateral quadriceps exercise (Rattey et al., 2006). Both of these suggest a strong central component of this phenomenon, although unlikely to be mediated by group III/IV afferents (Kennedy et al., 2015). However, we found no alteration of MUFR in the control limb from pre- to post- intervention, albeit when sampled at 25% MVC. Moreover, when estimating neural drive via the CST calculated from the firing times of MUs tracked pre to post, this also showed no alteration in the control limb. This limits the plausibility of the previous hypothesis of diminished neural drive as a causal factor to contralateral decrements. Although sampled in submaximal contractions in the current study, these were normalised to baseline MVC and therefore represent a higher *relative* force value but cannot be extrapolated across the higher recruitment range.

Non-local muscle fatigue may also be mediated by the circulation of metabolic by-products generated within the fatigued muscles (Halperin et al., 2015). The accumulation of metabolites such as potassium, hydrogen, heat-shock proteins, and blood lactate (Behm et al., 2021, Halperin et al., 2015) in the exercised muscle can migrate and exert non-local fatigue effects that impair enzymatic activity, muscle contractile dynamics, and action potential transmission (Cady et al., 1989, Kent-Braun, 1999). Additionally, repeated muscle activation is known to elevate extracellular potassium levels (Sejersted and Sjøgaard, 2000), which may migrate into non-exercise muscles. This repeated muscle activation alters the electrochemical gradients for potassium, which could contribute to diminished force production (Juel, 1986) and reduced excitability of the muscle, thereby contributing to performance fatigue (Nordsborg et al., 2003).

The current study found no change in MUFR variability at 25% MVC in either leg, which corresponds to previous studies with eccentric and concentric fatigue protocols (Jones et al., 2023). Additionally, we found no changes in MU recruitment and derecruitment thresholds in either leg, including of MUs tracked across the intervention. However, previous studies have shown increased (Farina et al., 2009) and decreased (Contessa et al., 2016) recruitment thresholds. These disparities in findings could be attributed to variations in task, muscle, and age groups considered.

The concurrent use of iEMG enabled a more detailed assessment of peripheral MU characteristics, and we found no RE-induced changes in MU properties in the exercise or contralateral control leg. The MUP negative peak ratio quantifies the relationship between the rise and fall slopes of the negative peak of the MUP template and may be sensitive to metabolite accumulation. In healthy young, this was altered following concentric fatigue protocols (lasting ∼1 hour), but not eccentric (Jones et al., 2023). These discrepancies could be primarily attributed to the variations in exercise protocols, particularly exercise duration. The NF jiggle is an indicator of NMJ transmission instability, which shows the variability of temporal dispersion across MU action potential (Piasecki et al., 2021) and is mainly related to the activity of muscle fibres close to the detection surface of the concentric needle electrode (within ∼350 µm) (Stashuk, 1999). Hence, the contributions of distant fibres to NF MUPs are minimal, which reduces the contamination from other MUs. However, although eccentric and concentric exercise protocols in healthy young individuals have been shown to increase NF jiggle (Jones et al., 2023), our resistance exercise protocol resulted in no change in NF jiggle in either leg. Again, it is probable these differences are explained by the shorter duration of the RE in the current study.

### Limitations

Firstly, the current study focused exclusively on immediate neuromuscular adaptations following a single bout of unilateral resistance exercise-induced performance fatigue. Therefore, these findings may not generalize to other types of intervention, such as those targeting endurance adaptations. Secondly, the current study reported MU data recruited at lower contraction levels (25% MVC) and therefore, these results may not be applicable for MUs with higher recruitment thresholds which are being recruited at higher force levels. Finally, this study did not include a healthy young cohort, so it was not possible to infer any age-related alterations in neuromuscular adaptations.

### Conclusions

The current study demonstrates that a single bout of unilateral knee extensor resistance exercise performed to fatigue is sufficient to elicit a bilateral reduction in maximal forces with neural adaptations occurring only in the exercise limb. These findings indicate that resistance exercise-induced muscle fatigue results in immediate central and peripheral alterations in the exercise limb. While the exact mechanism accounting for the force decline in the non-exercise limb remains uncertain, it is mainly suggestive of adaptations at systemic and/or central levels. These findings are important in understanding neurophysiological mechanisms following resistance exercise-based performance fatigue and may have potential implications for clinical settings including those often experienced by older adults (e.g. single-leg fracture, post-injury immobilisation, and stroke).

## Notes

### Competing Interest Statement

The authors have declared no competing interest.

## REFERENCES

Allen, D. G., Lamb, G. D. & Westerblad, H. 2008. Skeletal muscle fatigue: cellular mechanisms. Physiological reviews, 88, 287–332.

Altheyab, A., Alqurashi, H., England, T. J., Phillips, B. E. & Piasecki, M. 2024. Cross-education of lower limb muscle strength following resistance exercise training in males and females: A systematic review and meta-analysis. Experimental Physiology, 1–23.

Barss, T. S., Pearcey, G. E. & Zehr, E. P. 2016. Focus: The aging brain: Cross-education of strength and skill: An old idea with applications in the aging nervous system. The Yale journal of biology and medicine, 89, 81.

Bates, D., Mächler, M., Bolker, B. & Walker, S. 2015. Fitting Linear Mixed-Effects Models Using lme4. Journal of Statistical Software, 67, 1–48.

Behm, D. G., Alizadeh, S., Hadjizedah Anvar, S., Hanlon, C., Ramsay, E., Mahmoud, M. M. I., Whitten, J., Fisher, J. P., Prieske, O., Chaabene, H., Granacher, U. & Steele, J. 2021. Non-local Muscle Fatigue Effects on Muscle Strength, Power, and Endurance in Healthy Individuals: A Systematic Review with Meta-analysis. Sports Medicine, 51, 1893–1907.

Borg, E. & Kaijser, L. 2006. A comparison between three rating scales for perceived exertion and two different work tests. Scandinavian journal of medicine & science in sports, 16, 57–69.

Botter, A., Oprandi, G., Lanfranco, F., Allasia, S., Maffiuletti, N. A. & Minetto, M. A. 2011. Atlas of the muscle motor points for the lower limb: implications for electrical stimulation procedures and electrode positioning. European Journal of Applied Physiology, 111, 2461–2471.

Calvert, G. H. & Carson, R. G. 2022. Neural mechanisms mediating cross education: With additional considerations for the ageing brain. Neuroscience & Biobehavioral Reviews, 132, 260–288.

Contessa, P., De Luca, C. J. & Kline, J. C. 2016. The compensatory interaction between motor unit firing behavior and muscle force during fatigue. Journal of neurophysiology, 116, 1579–1585.

Debold, E. P. & Westerblad, H. 2024. New insights into the cellular and molecular mechanisms of skeletal muscle fatigue: the Marion J. Siegman Award Lectureships. American journal of physiology., 327, C946–C958.

Dehno, N. S., Kamali, F., Shariat, A. & Jaberzadeh, S. 2021. Unilateral strength training of the less affected hand improves cortical excitability and clinical outcomes in patients with subacute stroke: A randomized controlled trial. Archives of Physical Medicine and Rehabilitation, 102, 914–924.

Del Vecchio, A., Holobar, A., Falla, D., Felici, F., Enoka, R. M. & Farina, D. 2020. Tutorial: Analysis of motor unit discharge characteristics from high-density surface Emg signals. Journal of Electromyography and Kinesiology, 53, 102426.

Doix, A.-C. M., Lefèvre, F. & Colson, S. S. 2013. Time Course of the Cross-Over Effect of Fatigue on the Contralateral Muscle after Unilateral Exercise. PLos One, 8, e64910.

Dragert, K. & Zehr, E. P. 2013. High-intensity unilateral dorsiflexor resistance training results in bilateral neuromuscular plasticity after stroke. Experimental brain research, 225, 93–104.

Enoka, R. M. & Duchateau, J. 2016. Translating Fatigue to Human Performance. Medicine and science in sports and exercise, 48, 2228–2238.

Farina, D., Holobar, A., Gazzoni, M., Zazula, D., Merletti, R. & Enoka, R. M. 2009. Adjustments differ among low-threshold motor units during intermittent, isometric contractions. Journal of neurophysiology, 101, 350–359.

Francic, A. & Holobar, A. 2021. On the Reuse of Motor Unit Filters in High Density Surface Electromyograms Recorded at Different Contraction Levels. Ieee Access, 9, 115227–115236.

Gabriel, D. A., Kamen, G. & Frost, G. 2006. Neural adaptations to resistive exercise: mechanisms and recommendations for training practices. Sports medicine, 36, 133–149.

Guo, Y., Jones, E. J., Smart, T. F., Altheyab, A., Gamage, N., Stashuk, D. W., Piasecki, J., Phillips, B. E., Atherton, P. J. & Piasecki, M. 2024. Sex disparities of human neuromuscular decline in older humans. The Journal of Physiology, 603 (1), 151–165.

Halperin, I., Chapman, D. W. & Behm, D. G. 2015. Non-local muscle fatigue: effects and possible mechanisms. European Journal of Applied Physiology, 115, 2031–2048.

Holobar, A. & Zazula, D. 2007. Multichannel Blind Source Separation Using Convolution Kernel Compensation. Ieee Transactions on Signal Processing, 55, 4487–4496.

Inns, T. B., Bass, J. J., Hardy, E. J. O., Wilkinson, D. J., Stashuk, D. W., Atherton, P. J., Phillips, B. E. & Piasecki, M. 2022. Motor unit dysregulation following 15 days of unilateral lower limb immobilisation. The Journal of Physiology, 600, 4753–4769.

Jones, E. J., Guo, Y., Martinez-Valdes, E., Negro, F., Stashuk, D. W., Atherton, P. J., Phillips, B. E. & Piasecki, M. 2023. Acute adaptation of central and peripheral motor unit features to exercise-induced fatigue differs with concentric and eccentric loading. Experimental Physiology, 108, 827–837.

Jones, E. J., Piasecki, J., Ireland, A., Stashuk, D. W., Atherton, P. J., Phillips, B. E., Mcphee, J. S. & Piasecki, M. 2021. Lifelong exercise is associated with more homogeneous motor unit potential features across deep and superficial areas of vastus lateralis. GeroScience, 43, 1555–1565.

Juel, C. 1986. Potassium and sodium shifts during in vitro isometric muscle contraction, and the time course of the ion-gradient recovery. Pflügers Archiv, 406, 458–463.

Kawamoto, J.-E., Aboodarda, S. J. & Behm, D. G. 2014. Effect of differing intensities of fatiguing dynamic contractions on contralateral homologous muscle performance. Journal of sports science & medicine, 13, 836.

Kennedy, D. S., Fitzpatrick, S. C., Gandevia, S. C. & Taylor, J. L. 2015. Fatigue-related firing of muscle nociceptors reduces voluntary activation of ipsilateral but not contralateral lower limb muscles. Journal of applied physiology, 118, 408–418.

Lenth, R. & Lenth, M. R. 2018. Package ‘lsmeans’. The American Statistician, 34, 216–221.

Manca, A., Hortobágyi, T., Carroll, T., Enoka, R., Farthing, J., Gandevia, S., Kidgell, D., Taylor, J. L. & Deriu, F. 2021. Contralateral effects of unilateral strength and skill training: modified Delphi consensus to establish key aspects of cross-education. Sports Medicine, 51, 11–20.

Marathamuthu, S., Selvanayagam, V. S. & Yusof, A. 2022. Contralateral effects of eccentric exercise and Doms of the plantar flexors: evidence of central involvement. Research Quarterly for Exercise and Sport, 93, 240–249.

Martin, P. G. & Rattey, J. 2007. Central fatigue explains sex differences in muscle fatigue and contralateral cross-over effects of maximal contractions. Pflügers Archiv - European Journal of Physiology, 454, 957–969.

Martinez-Valdes, E., Negro, F., Botter, A., Pincheira, P. A., Cerone, G. L., Falla, D., Lichtwark, G. A. & Cresswell, A. G. 2022. Modulations in motor unit discharge are related to changes in fascicle length during isometric contractions. Journal of Applied Physiology, 133, 1136–1148.

Mcnair, P. J., Colvin, M. & Reid, D. 2011. Predicting maximal strength of quadriceps from submaximal performance in individuals with knee joint osteoarthritis. Arthritis care & research, 63, 216–222.

Mcphee, J. S., French, D. P., Jackson, D., Nazroo, J., Pendleton, N. & Degens, H. 2016. Physical activity in older age: perspectives for healthy ageing and frailty. Biogerontology, 17, 567–580.

Nordsborg, N., Mohr, M., Pedersen, L. D., Nielsen, J. J., Langberg, H. & Bangsbo, J. 2003. Muscle interstitial potassium kinetics during intense exhaustive exercise: effect of previous arm exercise. American Journal of Physiology-Regulatory, Integrative and Comparative Physiology, 285, R143–R148.

Parsaei, H., Nezhad, F. J., Stashuk, D. W. & Hamilton-Wright, A. Validation of motor unit potential trains using motor unit firing pattern information. 2009 Annual International Conference of the Ieee Engineering in Medicine and Biology Society, 2009. Ieee, 974–977.

Piasecki, M., Garnés-Camarena, O. & Stashuk, D. W. 2021. Near-fiber electromyography. Clinical Neurophysiology, 132, 1089–1104.

Piasecki, M., Ireland, A., Piasecki, J., Stashuk, D. W., Swiecicka, A., Rutter, M. K., Jones, D. A. & Mcphee, J. S. 2018. Failure to expand the motor unit size to compensate for declining motor unit numbers distinguishes sarcopenic from non-sarcopenic older men. The Journal of Physiology, 596, 1627–1637.

Rattey, J., Martin, P. G., Kay, D., Cannon, J. & Marino, F. E. 2006. Contralateral muscle fatigue in human quadriceps muscle: evidence for a centrally mediated fatigue response and cross-over effect. Pflügers Archiv - European Journal of Physiology, 452, 199–207.

Ruddy, K. L. & Carson, R. G. 2013. Neural pathways mediating cross education of motor function. Frontiers in human neuroscience, 7, 397.

Sejersted, O. M. & Sjøgaard, G. 2000. Dynamics and consequences of potassium shifts in skeletal muscle and heart during exercise. Physiological reviews, 80, 1411–1481.

Seo, D.-I., Kim, E., Fahs, C. A., Rossow, L., Young, K., Ferguson, S. L., Thiebaud, R., Sherk, V. D., Loenneke, J. P. & Kim, D. 2012. Reliability of the one-repetition maximum test based on muscle group and gender. Journal of sports science & medicine, 11, 221.

Singh, N. B., Arampatzis, A., Duda, G., Heller, M. O. & Taylor, W. R. 2010. Effect of fatigue on force fluctuations in knee extensors in young adults. Philosophical Transactions of the Royal Society A: Mathematical, Physical and Engineering Sciences, 368, 2783–2798.

Stashuk, D. 2001. Emg signal decomposition: how can it be accomplished and used? Journal of Electromyography and Kinesiology, 11, 151–173.

Stashuk, D. W. 1999. Detecting single fiber contributions to motor unit action potentials. Muscle & Nerve: Official Journal of the American Association of Electrodiagnostic Medicine, 22, 218–229.

Wilkinson, D. J., Piasecki, M. & Atherton, P. 2018. The age-related loss of skeletal muscle mass and function: Measurement and physiology of muscle fibre atrophy and muscle fibre loss in humans. Ageing research reviews, 47, 123–132.

